# Quantifying glioblastoma drug response dynamics incorporating resistance and blood brain barrier penetrance from experimental data

**DOI:** 10.1101/822585

**Authors:** Susan Christine Massey, Javier C. Urcuyo, Bianca Maria Marin, Jann N. Sarkaria, Kristin R. Swanson

## Abstract

Many drugs investigated for the treatment of glioblastoma (GBM) have had poor clinical outcomes, as their efficacy is dependent on adequate delivery to sensitive tumor cell populations, which is limited by the blood-brain barrier (BBB). Further complicating evaluation of therapeutic efficacy, tumors can become resistant to anti-cancer drugs, and it can be difficult to gauge the extent to which BBB limitations and resistance each contribute to a drug’s failure. To address this question, we developed a minimal mathematical model to characterize these elements of overall drug response, informed by time-series bioluminescence imaging data from a treated patient-derived xenograft (PDX) experimental model. By fitting this mathematical model to a preliminary dataset in a series of nonlinear regression steps, we estimated parameter values for individual PDX subjects that correspond to the dynamics seen in experimental data. Using these estimates, we performed a parameter sensitivity analysis using Latin hypercube sampling and partial rank correlation coefficients. Results from this analysis combined with simulation results suggest that BBB permeability may play a slightly larger role in therapeutic efficacy than drug resistance. Our model and fitting technique to estimate parameters from data may be a useful tool in aiding further exploration of these challenges in future studies of drug efficacy with larger datasets.

## 1 Introduction

Glioblastoma (GBM) is an aggressive primary brain cancer that is notoriously difficult to treat due to its diffuse infiltration into surrounding normal-appearing brain [10]. These diffusely invading GBM cells cannot be completely resected surgically [2], and are difficult to target with radiation therapy while sparing normal brain [5]. As a result, clinicians rely on chemotherapy to treat the full extent of the tumor. However, chemotherapeutic efficacy can be limited in two main ways: there may be insufficient delivery across the blood–brain barrier (BBB), and the tumor may develop resistance to therapy.

The BBB acts to keep pathogens and many toxins out of the sensitive brain tissue. Angiogenesis in dense tumor regions induces disruption of the BBB, potentially allowing chemotherapeutic drugs to “leak” into these tumor regions. Current dogma in neuro-oncology holds this as being largely sufficient to treat the tumor, but GBM cells invade beyond these regions into tissue where the BBB remains rather intact [19]. Further, tissue interstitial pressure and drug properties such as lipophilicity and polarity may influence the delivery of drugs across angiogenesis–induced BBB “leaks” [16]. Due to these factors, it remains unclear whether the delivery of BBB–impermeable antineoplastic agents reaches adequate concentrations throughout the tissue to provide the anticipated therapeutic effect.

Drug insensitivity or resistance is also a key suspect behind unsuccessful results treating GBM with molecularly–targeted therapies [7, 20]. GBMs frequently present with gene mutations or amplifications for a number of targets, such as epidermal growth factor receptor (EGFR), for which therapies already exist [3,4,8,15,18]. Due to the spatial heterogeneity of GBM, it is possible that these targets have been identified for a subpopulation of cells predominant in the dense core of the tumor where biopsies were taken, but are in fact less common in the invading portions of the tumor. This would result in a significant population of cells being treated with drugs they are not sensitive to, which could explain why trials with such targeted agents have failed [1, 6, 11, 17]. It is notable, however, that many of these drugs were not developed specifically for brain, but for cancers elsewhere in the body and subsequently tried in brain cancer, again raising the question of adequate delivery across the BBB.

“Specifically for EGFR or other kinase-targeted inhibitors, emergence of compensatory signaling pathways may lead to acquired resistance to therapy.”

In order to explore both the contributions of inadequate delivery of therapy across the BBB and drug resistance or insensitivity, we developed a minimal mathematical model based on preclinical experimental data. First, we describe model development based on this data and steps to estimate parameter regimes via data-fitting. Next, we explore the global model parameter sensitivity to understand how these parameters impact model outcomes. Finally, we run model simulations for the data–derived parameter regimes to assess the relative contributions of drug distribution and sensitivity, and discuss how it might be useful in assessing results from future experiments comparing different PDXs or different drugs. Overall, our model suggests that the influence of drug permeability may be more impactful than the degree of resistance for a given baseline sensitivity. Thus, in order to improve treatment outcomes, it is critical to determine predictors of drug distribution in individual patients’ tumors and surrounding brain tissue to ensure invading tumor cells are adequately exposed to the therapy.

## 2 Treatment Exposure and Sensitivity Model

Our ordinary differential equation (ODE) model of tumor growth and treatment response accounts for both variable treatment exposure and differential sensitivity to treatment by different tumor subpopulations. Development of this model was informed by experimental observations, which were also used to determine relevant parameter regimes for running simuliations.

### 2.1 Experimental Data

The form of our model was based on experimental data testing an EGFR-targeted antibody drug conjugate (ADC) [14]. These experiments were performed using a patient–derived xenograft (PDX) model of GBM, in which patient-derived GBM cells are implanted in rodents [9, 13]. The growth of these preclinical tumors was monitored via bioluminescent imaging (BLI, Figure 1). Since BLI flux is *linearly* correlated with tumor cell number [12], this provided us with a close approximation of tumor cell populations across time. Importantly, the PDX tumors were implanted both in the flank, where there is no BBB, and in the brain, which allowed us to compare treatment effect in tumors with and without BBB impediments to drug distribution. Moreover, there were control groups in both sites (flank and intracranial) that received no treatment.

**Figure 1:**
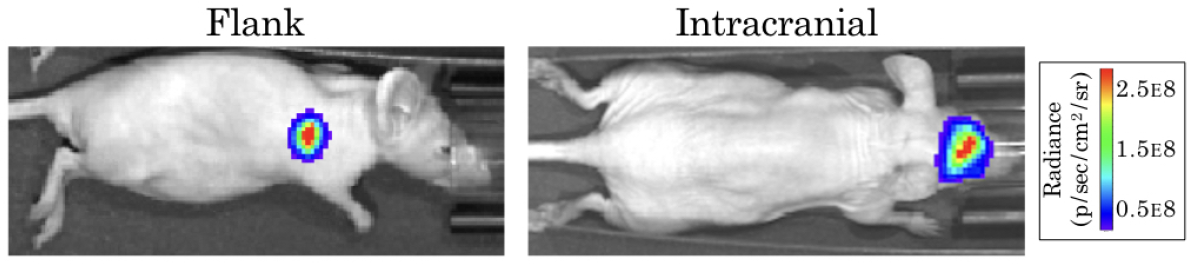
Example bioluminescence images for patient-derived xenografts. Colors represent the BLI radiance in photons per second per area in cm squared.., which is related to the BLI flux (measured in photons per second) for the total area.

In the untreated groups, BLI flux increased exponentially over time, indicating exponential growth in the total tumor cell population. Treated groups showed similar exponential growth in the absence of treatment at early time points, followed by a precipitous decline with the initiation of therapy (begun once tumors reached a set volume). This period of decline was relatively short–lived, and after that BLI flux increased exponentially again, albeit at a slower rate. The precipitous decline in experimental tumor volume followed by a trajectory of slower regrowth than that prior to therapy suggested the presence of two distinct tumor cell populations: one that responded to therapy (sensitive, denoted *s*), and one that continued to grow in spite of therapy (resistant, denoted *r*). These cell populations are assumed to proliferate at the same rate, *ρ*, differing only in their responses to treatment. The ADC therapy (denoted *A*) was delivered every seven days at dose *A*_dose_, and decays at rate *λ*. In response to therapy, which achieves a fraction *γ* of the dosed concentration in the tumor tissue, cell death is induced in each population at rates (*μ*_*s,r*_), respectively. These dynamics are schematized in Figure 2.

**Figure 2:**
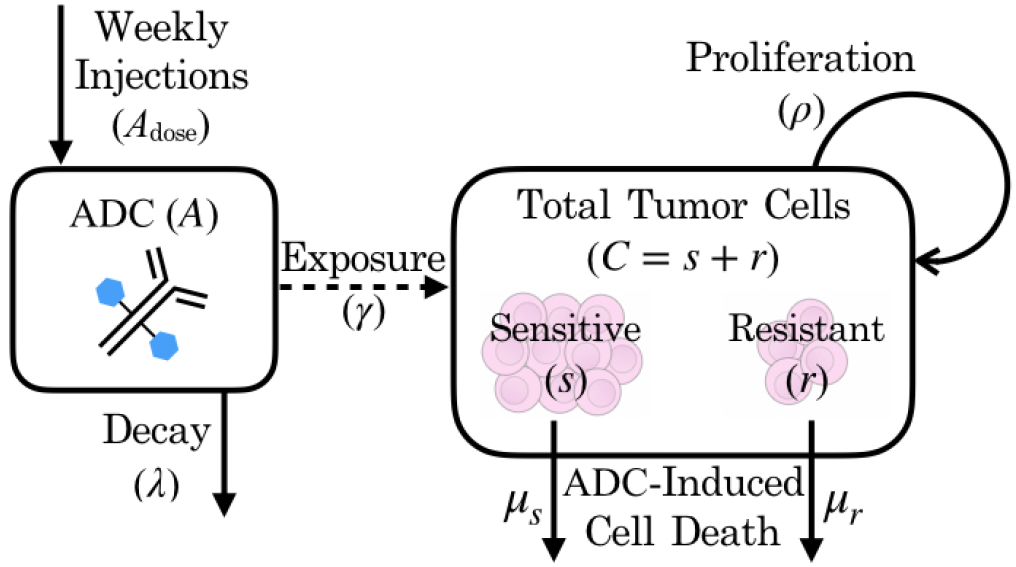
Schematic of patient-derived xenograft response to an antibody drug conjugate, including key variables and parameters of the mathematical model.

### 2.2 Model Equations

Our model consists of three coupled ordinary differential equations describing the dynamics of both cell populations (*s*, *r*) and the ADC (*A*):

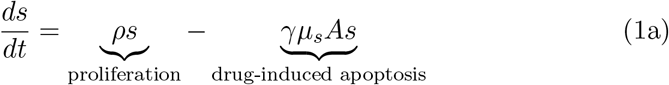

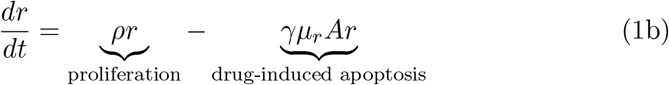

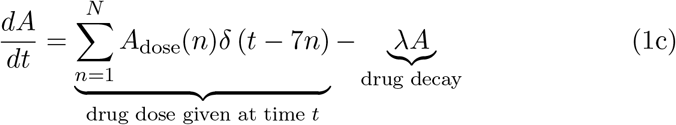

where parameters and their definitions are outlined in Table 1, and their derivations can be found in Section 2.3.

**Table 1:** Model Parameter Definitions and Values. Parameter value ranges were estimated through fitting the model to experimental data or parameters were confined to a value range by their theoretical meaning, except in the case of ADC parameters, as described in Section 2.3.

| Symbol | Definition | Value Range | Units |
| --- | --- | --- | --- |
| $\rho$ | cellular proliferation rate | 0.2 to 0.5 | day<sup>-1</sup> |
| $\mu_s$ | ADC-mediated sensitive cell kill rate | 1 to 10 | mg<sup>-1</sup> day<sup>-1</sup> |
| $\mu_r$ | ADC-mediated resistant cell kill rate | $z\mu_s$ | mg<sup>-1</sup> day<sup>-1</sup> |
| $q$ | resistant proportion of implanted cells | 0 to 1 | unitless |
| $z$ | sensitivity factor between $\mu_s$ & $\mu_r$ | 0 to 1 | unitless |
| $\lambda$ | rate of ADC decay | $\ln(2)/7$ | day<sup>-1</sup> |
| $\gamma$ | proportion of tumor exposed | 0 to 1 | unitless |
| $A_{\text{dose}}$ | ADC given in a single dose | 0.1 | mg |

In the absence of the ADC, both sensitive and resistant tumor populations grow exponentially, at proliferation rate *ρ*. However, the two populations differ in sensitivity to the ADC, *A*, which is captured by the drug-induced apoptosis rates *μ*_*s*_ and *μ*_*r*_ (for sensitive and resistant populations, respectively). The terms for tumor cell death due to ADC are further modified by factor *γ*, which represents the proportion of cells exposed to ADC. We assume that the ADC is readily distributed to flank PDXs such that tumor cell exposure is high (*γ* = 1), but that the BBB limits this distribution for intracranial PDXs (0 ≤ *γ* ≤ 1). In order to capture the ADC dynamics, we let *A*_dose_(*n*) represent the *n*th dose, with doses administered every seven days, as noted by the dirac delta function *δ*(*t* − 7*n*). The ADC then decays at rate *λ*.

This model can be solved analytically, as shown in Appendix A. For simplification, at any given time *t*, *C*(*t*) represents the total number of cells, calculated by the sum of sensitive *s*(*t*) and resistant *r*(*t*) cells. This total cell number was used in Section 2.3 for comparing with bioluminescence imaging data, which shows the total tumor cell population. The initial resistant proportion of total implanted cells is denoted by *q* = *r*_0_/*C*_0_. Similarly, the extent to which resistant cells *r* are less sensitive to the agent than the sensitive cells *s* is denoted by the ratio *z* = *μ*_*r*_/*μ*_*s*_, which is bounded between 0 and 1 to ensure that *μ_r_* is a fraction of *μ*_*s*_ in the regression-based parameterization in Section 2.3. With these notational changes, we can then write the analytical solution (derived in Appendix A) as

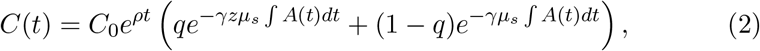

where

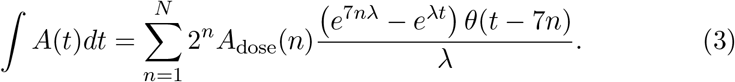

Utilizing this solution (2), the model can be parameterized through comparison of simulations to temporal BLI data.

### 2.3 Data-Based Parameter Estimation

Most model parameters were unknown, with the exception of ADC-specific parameters: the timing of dose administration and dose amounts (*A*_dose_(*n*)), as well as the half-life of the drug, which allowed us to solve for the drug decay rate (*λ*). Dose amounts were adjusted for the weight of each animal (5 mg/kg), so we applied the average initial animal weight of 20 g to obtain the constant ADC dose, *A*_dose_ = 0.1 mg used in simulations. All of the remaining model parameters were determined through several iterations of fitting the model via least squares regression to preliminary BLI data from an experiment. The various arms of the experiment included untreated and treated groups of subjects, as well as flank and intracranial tumor sites to separate out BBB influences. By fitting the model to these various sub-groups, we are able to identify and estimate each of the parameters, as described below.

#### Step 1 Fit to untreated data to estimate growth rate, *ρ*

When fitting the model to untreated data, since the ADC is not injected (*A* = 0), the model’s treatment components zero out and only an exponential growth function remains: *C*(*t*) = *C*_0_*e*^*ρt*^. Fitting this model function to untreated data via least squares regression with the `lsqcurvefit` function in MATLAB® (MATLAB Release 2018b, The MathWorks, Inc., Natick, Massachusetts, United States) (Figure 6), we were able to obtain estimates of the tumor proliferation rate *ρ* and the number of viable implanted PDX cells *C*_0_. (Note that while a consistent number of cells are initially implanted for each subject, *C*_0_ is in fact unique for each, as a variable number of cells die off, possibly due to an inability to establish themselves in the proper microenvironment for growth.) This yielded subject-specific values for *ρ* and *C*_0_, and the mean *ρ* was recorded as the net proliferation rate for the cells of the particular PDX line used in the experiments grown in either the flank or intracranial setting.

#### Step 2 Fit to treated flank data to estimate *μ*_*s*_, *z*, and *q*

Using the estimated net proliferation rate, *ρ*, from the previous step, we proceed to fit the treated data in the flank. We assume the flank-specific estimate of *ρ* remains the same in the treated case as untreated, since the microenvironment remains similar and any effects on growth can be encapsulated in the treatment effect term. Additionally, since the tumor was injected in the flank, there is no BBB effect to limit the proportion of tumor exposed to the ADC, such that the exposure parameter *γ* = 1. Pairing this with other known parameters (see Table 1), the only remaining unknown parameters are the cell death rates due to drug, *μ*_*s*_ and *μ*_*r*_, and proportion of implanted cells that are resistant, *q*. Using the definition *z* = *μ*_*r*_/*μ*_*s*_ and the analytical solution of the model (2), we can then apply a nonlinear least squares regression (again using lsqcurvefit) to fit subject-specific parameters for parameters *μ*_*s*_, *z*, and *q* (Figure 6).

#### Step 3 Fit to treated intracranial data to estimate *γ*, *z*, and *q*

Proceeding to fit the data from treated intracranial tumors, we apply the same approach to estimate parameters as in the flank, this time assuming that the intracranial-specific estimate of *ρ* from the untreated setting remains the same for the treated intracranial tumors due to a similar microenivronment. We further assume that the cell death rate due to therapy for the sensitive tumor subpopulation, *μ*_*s*_, is the same intracranially as in the flank setting and estimate parameter *γ*, the fraction of tumor exposed to therapy.

At the conclusion of these steps (summarized in Figure 3), all unknown model parameters had net and individual estimates (listed in Table 1). Using these values then allowed us to run simulations in a reasonable range of parameter values, as well as to perform a model sensitivity analysis to understand how variability in these values affect model outcomes.

**Figure 3:**
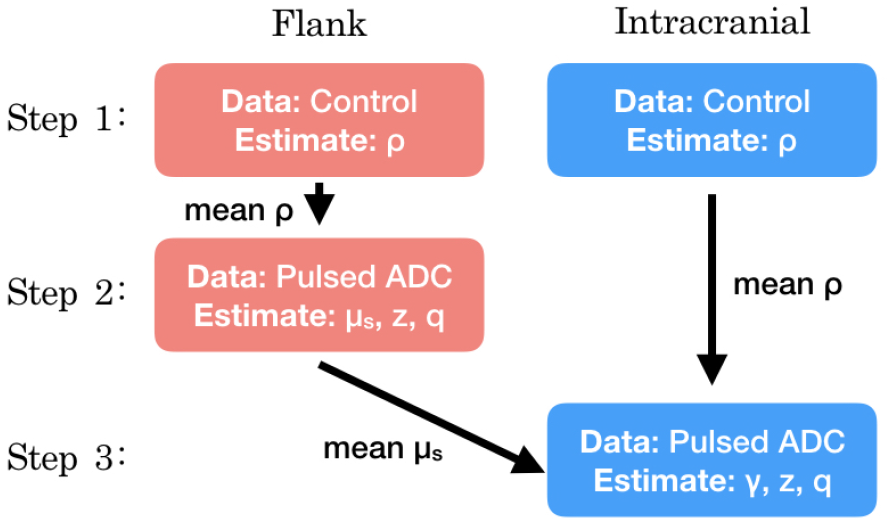
Summary of the series of steps used to estimate model parameter values through fitting the model to different experimental data sets.

## 3 Results

### 3.1 Parameter Sensitivity Analysis

Due to the uncertainty and variability in our parameter estimates, it was important to better characterize the effects of parameters on model results. To do this, we conducted a parameter sensitivity analysis via Latin hypercube sampling (LHS) and partial–rank correlation coefficients (PRCC). To perform the LHS analysis, we first drew 1000 equiprobable samples for each unknown parameter, including the initial condition *C*_0_, from a statistical distribution of values. These distributions were informed by our fits of the preliminary data when available; in the case of the unitless parameters, we assumed a uniform distribution on the interval [0, 1]. These samples were then randomly paired in a Latin hypercube scheme to run a series of 1000 Monte Carlo simulations. Using these simulation results, we then computed PRCCs between each parameter and two different model outcomes across all time points: the total number of tumor cells and the fraction of tumor that is resistant (Figure 4). Further details for about this method and the code files used are available on GitHub: https://github.com/scmassey/model-sensitivity-analysis.

**Figure 4:**
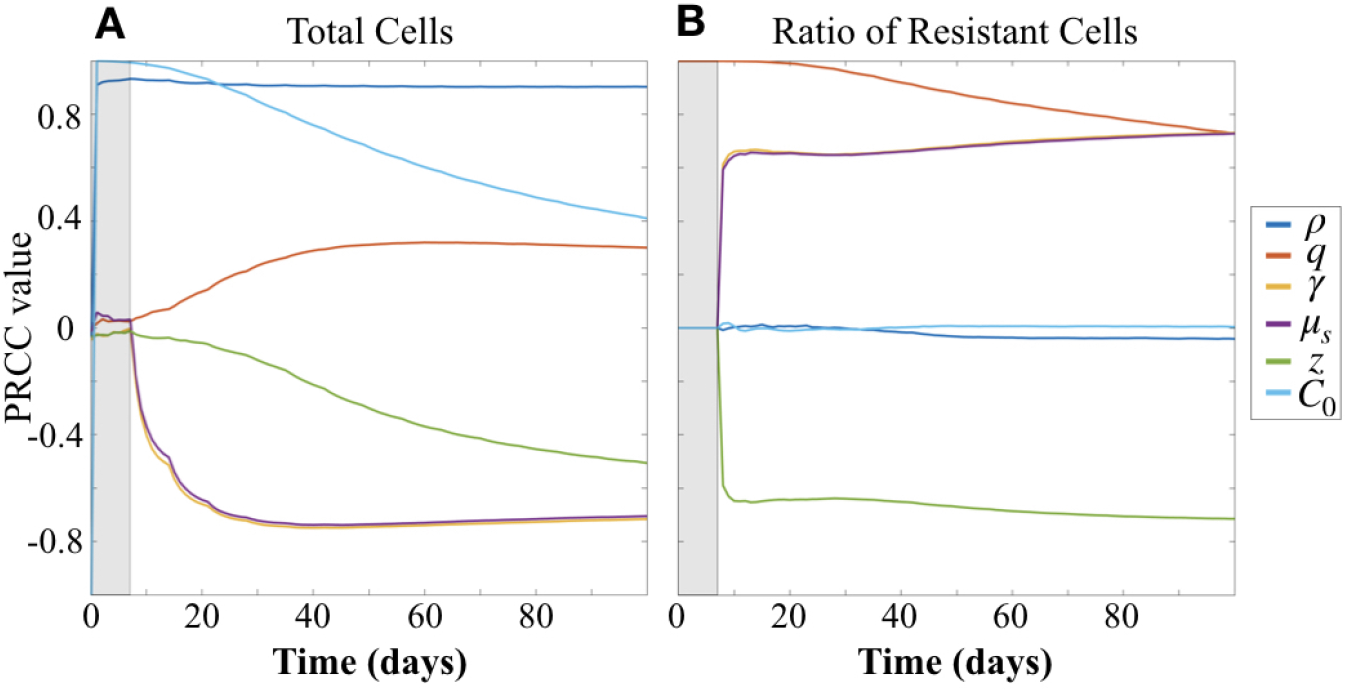
Partial rank correlation coefficients (PRCC) of parameters with respect to (A) tumor cells and (B) fraction of resistant cells (*r*/(*s* + *r*) = *r*/*C*), visualized across simulation time.

#### 3.1.1 Total tumor population depends most strongly on proliferation rate, followed by treatment response parameters

At early time points, particularly before the initiation of therapy, the tumor population is strongly positively correlated with both the initial number of cells implanted, *C*_0_ and proliferation rate *ρ* (Figure 4A). However by 30 days, or after approximately three doses of therapy, the population is strongly positively correlated with *ρ* and the effect of *C*_0_ begins to wane. At the same time, drug sensitivity of the *s* cell population, *μ*_*s*_, and exposure to drug, *γ* are strongly negatively correlated. Resistance factor *z*, which determines the fraction of drug sensitivity remaining in the *r* cell population, is also negatively correlated, but less strongly, and only reaches a PRCC value of −0.5 after 100 days. This suggests that resistance *z* is less impactful than the baseline sensitivity of the tumor population and the distribution of drug in the tissue.

As expected from our substitution of *μ*_*r*_ = *zμ*_*s*_, which results in the coefficient −*γzμ*_*s*_ in the term describing drug induced apoptosis for the equation describing the *r* population (1b), which mirrors that for the *s* population (1a), −*γμ*_*s*_, parameters *γ* and *μ*_*s*_ track together in the sensitivity analysis (overlapping lines in Figure 4). Thus, sensitivity analysis is unable to compare the differential impacts of these two parameters. However, we note that since we had preliminary data in both the treated flank as well as the treated intracranial PDX settings, we were able to obtain parameter estimates for these by keeping *γ* = 1 in the flank setting, and assuming that *μ*_*s*_ is the same intracranially as in flank.

#### 3.1.2 Resistant fraction of tumor driven by initial resistant proportion, followed by treatment response parameters

Prior to the initiation of therapy, only parameter *q*, the fraction of initially implanted cells that are resistant, is correlated with the proportion of total tumor that is resistant (Figure 4B). Once treatment is initialized, *q* remains highly positively correlated, and this correlation decreases slightly over time during the course of treatment.

Three other parameters show correlation with the fraction of tumor that is resistant following initiation of therapy, all of which involve drug response. Parameters *γ* and *μ*_*s*_, representing the degree of exposure to drug and baseline sensitivity of the cells, respectively, are both positively correlated and track together, while parameter *z*, representing the fraction of treatment sensitivity remaining in resistant cells, is negatively correlated with the fraction of tumor that is resistant. Further, the PRCC values do not vary over the time of the simulation after treatment is initiated and sustained. These correlations are consistent with expectations from the behavior of the system described by the model.

### 3.2 Simulation Results

To more fully explore the effect of parameters on model predicted outcomes, we ran simulations for varied values of the parameters relating to treatment response: *γ*, *μ*_*s*_, and *z*, or exposure to therapy, sensitivity to therapy, and degree of reduced sensitivity to therapy in resistant cells, respectively. Codes used to run simulations and plot the results may be found on GitHub: https://github.com/scmassey/treatment-exposure-sensitivity-model.

#### 3.2.1 Treatment exposure is especially important for tumors with lower baseline sensitivity

Comparing simulation results across a range of values for parameters *γ* and *z*, we see that *γ* plays a larger influence on total tumor cells for a given baseline sensitivity than does *z*. That is, our model suggests that exposure to therapy is more impactful on total tumor burden than the degree of resistance, which is consistent with the parameter sensitivity results of Section 3.1.1. The simulations also indicate that for higher sensitivity, treatment can be effective at lower levels of exposure (Figure 5, compare heatmaps for *μ*_*s*_ = 3 vs *μ*_*s*_ = 7).

**Figure 5:**
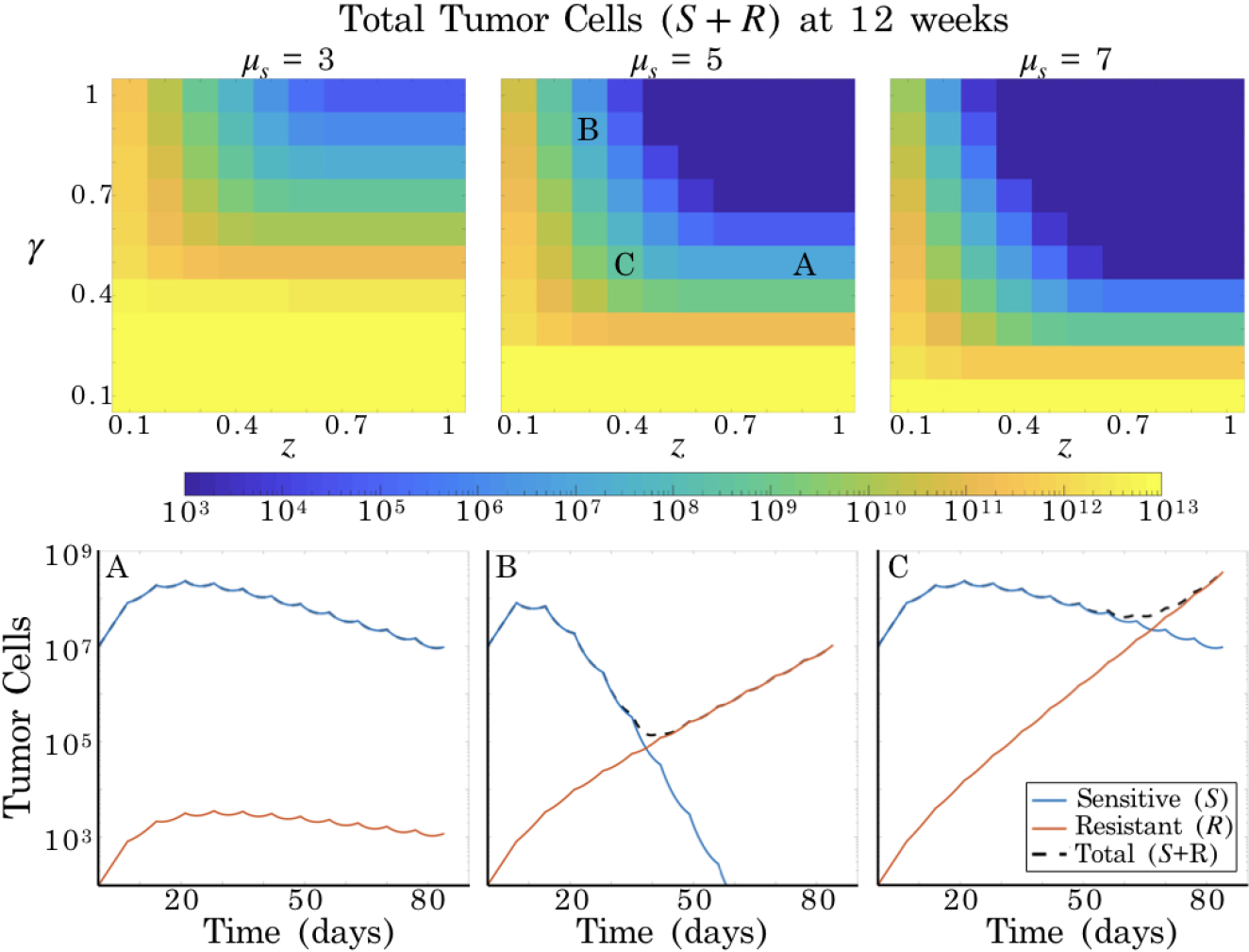
Simulation results for varied *γ* and *z*. Top row: heatmaps show total tumor cells at 84 days (12 weeks) post tumor initiation, across ten values each of *γ* and *z*, for three values of *μ*_*s*_. Recall that *μ*_*r*_ = *zμ*_*s*_, such that it is straightforward to compute the corresponding *μ*_*r*_ value for a given *z* value in each of the heatmaps. Bottom row: three particular simulations corresponding to the labels in the heatmap associated with *μ*_*s*_ = 5.

#### 3.2.2 Same total tumor burden, different fractions of sensitive vs resistant subpopulations

Further, we observed that there can be distinct differences in the dynamics of the individual cell populations underlying simulations that show the same resulting overall tumor cell population level (Figure 5A,B). Looking at long time scales (> 50 days)—in this case at 12 weeks or 84 days, the average survival time of the treated subjects—we are able to observe the effect of an extended time of treatment in the simulations. It is particularly notable that one simulation retains a large proportion of treatment sensitive cells (Figure 5A), while another is made up almost entirely of treatment resistant cells (Figure 5B). These two simulations share the same baseline sensitivity, *μ*_*s*_, and the same level of exposure, *γ*, differing only in the degree *z* to which the resistant subpopulation is insensitive to the ADC.

### 3.3 Parameter Estimation for comparing across subjects and experimental groups

While only performed for a small set of data, we note that our parameter estimation procedure may be useful for comparing results within and across experiments. In this small data set, the variation in growth curves between the five treated flank and intracranial subjects was associated with differences in estimated parameter values. (This data, with overlaid simulations using the fitted parameter estimates are shown in Figure 6.) Comparing these values within and across different treatments may provide clues as to the origins of heterogeneous trial results.

**Figure 6:**
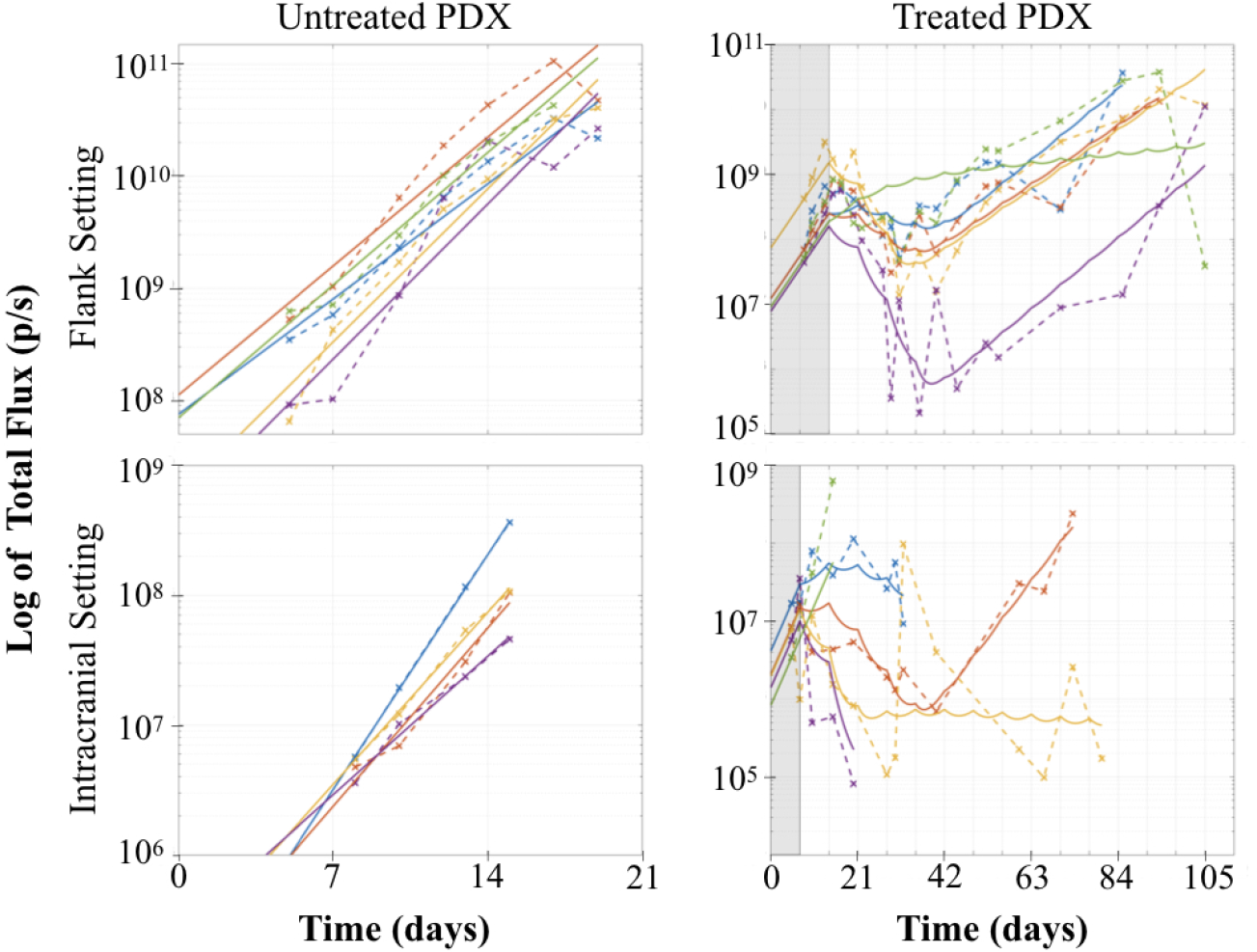
Tumor growth assessed by bioluminescence imaging flux (dashed lines), with model fits (solid lines). On the right panel, the shaded region indicates the time before treatment initiated. Note that the y-axis is log scaled.

## 4 Discussion

Our Treatment Exposure and Sensitivity model describes tumor growth and treatment response incorporating the effects of the BBB on tumor exposure to therapy as well as differential therapeutic sensitivity. It is important to note that tumor heterogeneity may actually provide for many subpopulations with varying levels sensitivity to therapy; we assumed that these cluster towards more or less sensitive, reducing these to two subpopulations. Additionally, the therapy may have an effect not only through induction of tumor cell apoptosis, but could also reduce tumor proliferation. Because our data does not provide for an ability to distinguish between these effects, both are encapsulated in the *μ*_*s*,*r*_ therapeutic effect parameters. This assumption allows us to use the estimates of *ρ* obtained through fitting the model to untreated flank and intracranial experimental groups in the treated flank and intracranial settings.

Parameter sensitivity analysis of the model highlighted the necessity of having the flank and intracranial treatment groups for practical identifiability in obtaining estimates for *γ* and *μ*_*s*_. This “tradeoff” between therapeutic sensitivity (*μ*_*s*_) and exposure (*γ*) was also observed in simulations and is expected given our model formulation. However, the emergence of this dynamic in the creation and parameter estimation of our model underscores that this relationship should be carefully considered in the design of experimental studies for new glioma therapies.

Sensitivity analysis also revealed that parameter *ρ* is the strongest positive influence on total cell population, as expected, and distinguished between the effect of reduced therapeutic sensitivity (*z*) and exposure to therapy (*γ*) in reducing the total tumor population. Not only is there a difference in the magnitude of correlation between parameters *γ* and *z* with total tumor burden, there is also a difference in the temporal dynamics of the change in these correlations over time. Further, most correlation coefficients between parameters and total tumor burden change most dramatically at earlier time points, and after approximately 50 days, change relatively little. This suggests that experiments conducted to examine the relationship between exposure and sensitivity to therapy should focus on collecting time course data more densely for the first 7 weeks as compared to longer times. It is also notable that the correlation coefficients between parameters and the resistant fraction of the tumor is quite stable over time. Simulations confirmed the greater impact of therapeutic exposure (*γ*) relative to reduction in sensitivity (*z*) foreshadowed by the sensitivity analysis. More importantly, they revealed that for a particular tumor burden at any *single* time, there can be several parameterizations that fit the data but correspond to different fractions of resistant cells. This indicates that time course data is essential for detecting differences in the contributions of *γ* and *z*.

Finally, because the model we have presented is minimal and reduces mechanisms down to a few key parameters, it has greater utility for fitting experimental data to estimate these parameters for individuals. Having these individual parameterizations is key to understanding the extent to which drug exposure and resistance each contributed to variations in outcome. We anticipate using this approach in the future for comparing experimental groups with additional PDX lines as well as other therapies, characterizing the differences in outcome results between those groups. This will enable us to determine whether there is any consistency in parameter values within groups, which may lead to improved understanding of patterns behind drug failures and identification of tumor characteristics that might suggest candidates who would benefit most from an emerging therapy.

## A Analytical Solution to Model

Here we present the derivation of the analytical solution for model system (1). Recall that the system consists of the following differential equations, which describe the growth of two tumor cell populations with differential drug sensitivity, *s* and *r*, and drug level, *A*:

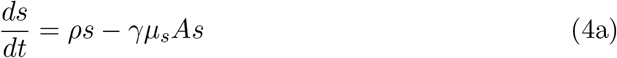

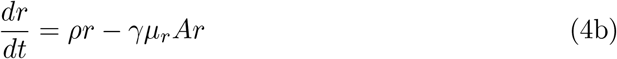

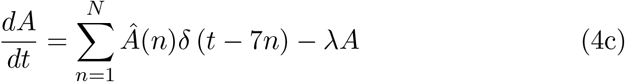

where *Â* is a vector containing the consecutive drug doses administered, such that *Â*(*n*) is the dose on the *n*th pulse (of which there are at most *N* drug pulses), and 7*n* denotes the time of the *n*th pulse (every 7 days). The parameters are given in Table 1 of the main text.

Because the equation for *A* in the system (4) is independent of *s* and *r*, we first solve (4c) for *A*. First, observe the following rearrangement:

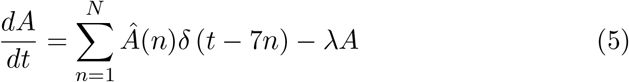

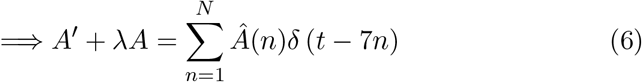

Notice that we can use the method of integrating factors, introducing a factor of *e*^*λt*^:

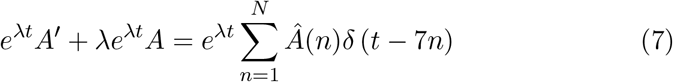

so that

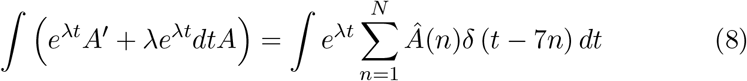

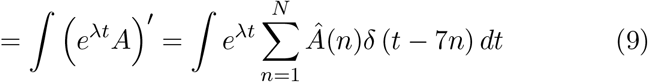

Integrating the left hand side, and moving the integral inside the sum on the right hand side:

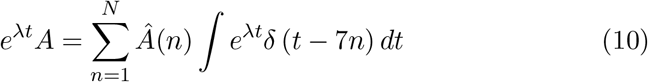

Then, integrating the right hand side, we have

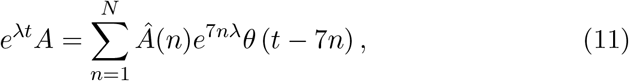

where *θ* is the heaviside function.

Now, since *λ* = ln(2)/7 (i.e., the half life is 7 days), we have the coefficient

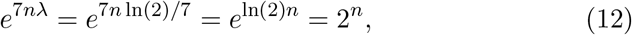

so the right hand side of (11) becomes

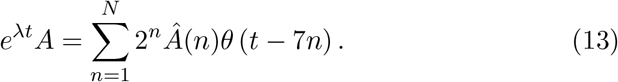

Rearranging, we have our solution for *A*:

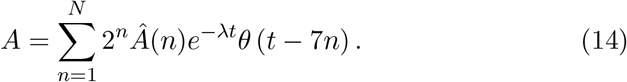

Now we solve for *s* and *r*, and use the fact that (4a) and (4b) differ only by parameter *μ*_*s*_ versus *μ*_*r*_ to obtain both simultaneously:

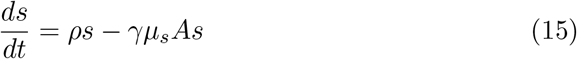

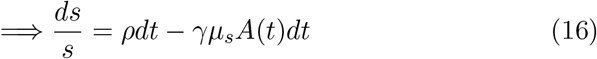

Integrating both sides:

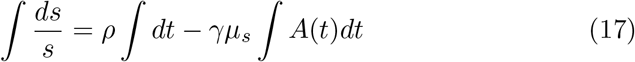

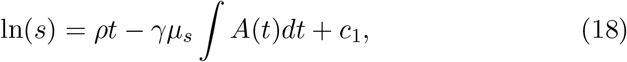

where *c*_1_ is a constant of integration to be determined from the initial condition.

Since the solution for *s* in (18) depends on *∫ A*(*t*)*dt*, we must integrate (14):

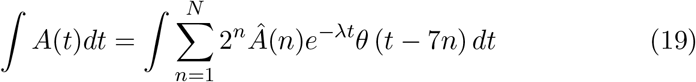

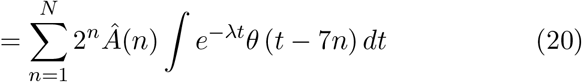

To use integration by parts, such that *∫ u*′*v* = *uv* − *uv*′, choose

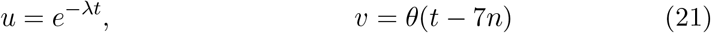

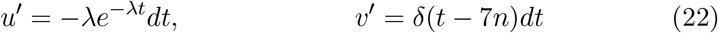

Then we have:

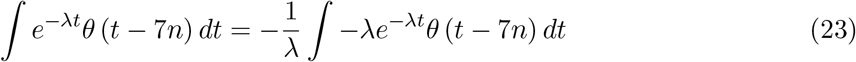

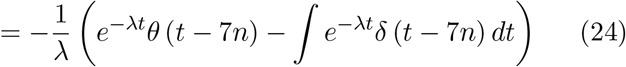

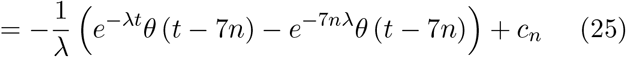

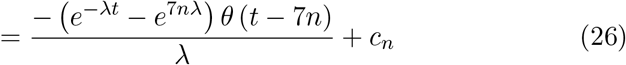

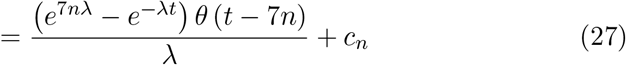

where *c*_*n*_ is a constant of integration. Thus, (20) becomes:

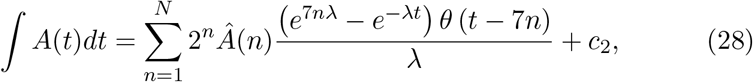

where *c*_2_ is the sum of all the *c*_*n*_.

Now we can insert (28) into (18), giving

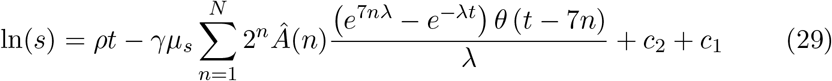

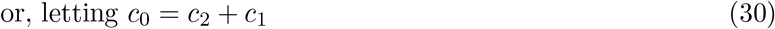

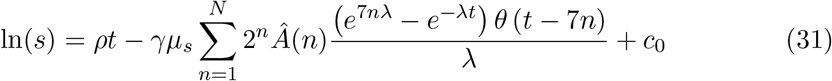

Solving for *s* then, we find

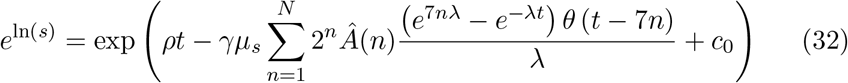

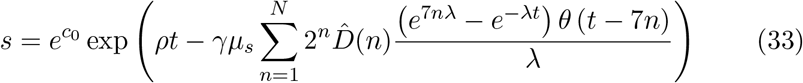

Denoting exp(*c*_0_) = *s*_0_, we arrive at our solution for *s*:

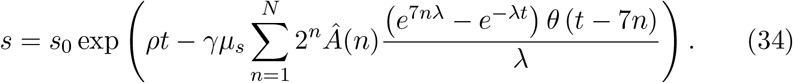

The solution for *r* is similar:

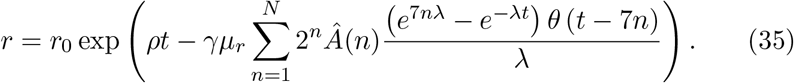

Now, we assume that the *s* cells are more sensitive to drug than the *r* cells, such that *μ*_*s*_ > *μ*_*r*_. Defining *z* to be the ratio *z* = *μ*_*r*_/*μ*_*s*_, we can substitute *μ*_*r*_ = *zμ*_*s*_, and in our fitting process, solve for this degree of differential sensitivity between the *s* and *r* cell populations.

Additionally, for fitting purposes, we can combine the cells into one total population: *C* = *s* + *r*. To do this, we write the initial condition *C*_0_ = *s*_0_ + *r*_0_, and let *s*_0_ = *qA*_0_ such that *q* represents the proportion of *C*_0_ that is made up of cells that are more drug–sensitive. This gives:

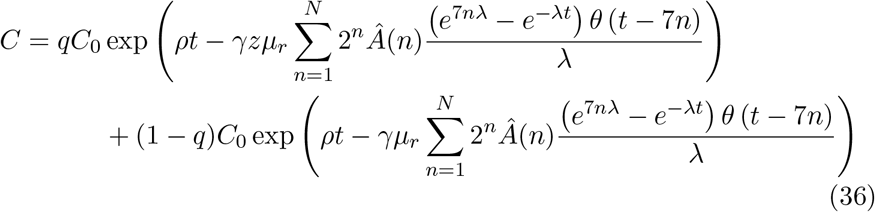

Because this is a lengthy expression, in the main text we write

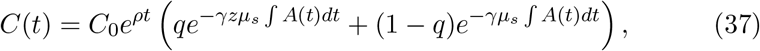

where *∫ Adt* denotes the summation in (20).

## References

[1] Abbvie provides update on depatuxizumab mafodotin (depatuxm), an investigational medicine for newly diagnosed glioblastoma, an aggressive form of brain cancer [Press Release]. https://news.abbvie.com/news/press-releases/abbvie-provides-update-on-depatuxizumab-mafodotin-depatux-m-an-investigational-medicine-for-newly-diagnosed-glioblastoma-an-aggressive-form-brain-cancer.htm (2019). Accessed: 2019-09-03

[2] Baldock, A.L., Ahn, S., Rockne, R., Johnston, S., Neal, M., Corwin, D., Clark-Swanson, K., Sterin, G., Trister, A.D., Malone, H., et al.: Patient-specific metrics of invasiveness reveal significant prognostic benefit of resection in a predictable subset of gliomas. PLoS One 9(10), e99,057 (2014)

[3] van den Bent, M., Gan, H.K., Lassman, A.B., Kumthekar, P., Merrell, R., Butowski, N., Lwin, Z., Mikkelsen, T., Nabors, L.B., Papadopoulos, K.P., et al.: Efficacy of depatuxizumab mafodotin (abt-414) monotherapy in patients with egfr-amplified, recurrent glioblastoma: results from a multi-center, international study. Cancer Chemotherapy and Pharmacology 80(6), 1209–1217 (2017). DOI 10.1007/s00280-017-3451-1. URL http://dx.doi.org/10.1007/s00280-017-3451-1

[4] Brennan, C.W., Verhaak, R.G., McKenna, A., Campos, B., Noushmehr, H., Salama, S.R., Zheng, S., Chakravarty, D., Sanborn, J.Z., Berman, S.H., Beroukhim, R., Bernard, B., Wu, C.J., Genovese, G., Shmulevich, I., Barnholtz-Sloan, J., Zou, L., Vegesna, R., Shukla, S.A., Ciriello, G., Yung, W., Zhang, W., Sougnez, C., Mikkelsen, T., Aldape, K., Bigner, D.D., Meir, E.G.V., Prados, M., Sloan, A., Black, K.L., Eschbacher, J., Finocchiaro, G., Friedman, W., Andrews, D.W., Guha, A., Iacocca, M., O’Neill, B.P., Foltz, G., Myers, J., Weisenberger, D.J., Penny, R., Kucherlapati, R., Perou, C.M., Hayes, D.N., Gibbs, R., Marra, M., Mills, G.B., Lander, E., Spellman, P., Wilson, R., Sander, C., Weinstein, J., Meyerson, M., Gabriel, S., Laird, P.W., Haussler, D., Getz, G., Chin, L., Benz, C., Barnholtz-Sloan, J., Barrett, W., Ostrom, Q., Wolinsky, Y., Black, K.L., Bose, B., Boulos, P.T., Boulos, M., Brown, J., Czerinski, C., Eppley, M., Iacocca, M., Kempista, T., Kitko, T., Koyfman, Y., Rabeno, B., Rastogi, P., Sugarman, M., Swanson, P., Yalamanchii, K., Otey, I.P., Liu, Y.S., Xiao, Y., Auman, J., Chen, P.C., Hadjipanayis, A., Lee, E., Lee, S., Park, P.J., Seidman, J., Yang, L., Kucherlapati, R., Kalkanis, S., Mikkelsen, T., Poisson, L.M., Raghunathan, A., Scarpace, L., Bernard, B., Bressler, R., Eakin, A., Iype, L., Kreisberg, R.B., Leinonen, K., Reynolds, S., Rovira, H., Thorsson, V., Shmulevich, I., Annala, M.J., Penny, R., Paulauskis, J., Curley, E., Hatfield, M., Mallery, D., Morris, S., Shelton, T., Shelton, C., Sherman, M., Yena, P., Cuppini, L., DiMeco, F., Eoli, M., Finocchiaro, G., Maderna, E., Pollo, B., Saini, M., Balu, S., Hoadley, K.A., Li, L., Miller, C.R., Shi, Y., Topal, M.D., Wu, J., Dunn, G., Giannini, C., O’Neill, B.P., Aksoy, B.A., Antipin, Y., Borsu, L., Berman, S.H., Brennan, C.W., Cerami, E., Chakravarty, D., Ciriello, G., Gao, J., Gross, B., Jacobsen, A., Ladanyi, M., Lash, A., Liang, Y., Reva, B., Sander, C., Schultz, N., Shen, R., Socci, N.D., Viale, A., Ferguson, M.L., Chen, Q.R., Demchok, J.A., Dillon, L.A., Shaw, K.R.M., Sheth, M., Tarnuzzer, R., Wang, Z., Yang, L., Davidsen, T., Guyer, M.S., Ozenberger, B.A., Sofia, H.J., Bergsten, J., Eckman, J., Harr, J., Myers, J., Smith, C., Tucker, K., Winemiller, C., Zach, L.A., Ljubimova, J.Y., Eley, G., Ayala, B., Jensen, M.A., Kahn, A., Pihl, T.D., Pot, D.A., Wan, Y., Eschbacher, J., Foltz, G., Hansen, N., Hothi, P., Lin, B., Shah, N., geun Yoon, J., Lau, C., Berens, M., Ardlie, K., Beroukhim, R., Carter, S.L., Cherniack, A.D., Noble, M., Cho, J., Cibulskis, K., DiCara, D., Frazer, S., Gabriel, S.B., Gehlenborg, N., Gentry, J., Heiman, D., Kim, J., Jing, R., Lander, E.S., Lawrence, M., Lin, P., Mallard, W., Meyerson, M., Onofrio, R.C., Saksena, G., Schumacher, S., Sougnez, C., Stojanov, P., Tabak, B., Voet, D., Zhang, H., Zou, L., Getz, G., Dees, N.N., Ding, L., Fulton, L.L., Fulton, R.S., Kanchi, K.L., Mardis, E.R., Wilson, R.K., Baylin, S.B., Andrews, D.W., Harshyne, L., Cohen, M.L., Devine, K., Sloan, A.E., VandenBerg, S.R., Berger, M.S., Prados, M., Carlin, D., Craft, B., Ellrott, K., Goldman, M., Goldstein, T., Grifford, M., Haussler, D., Ma, S., Ng, S., Salama, S.R., Sanborn, J.Z., Stuart, J., Swatloski, T., Waltman, P., Zhu, J., Foss, R., Frentzen, B., Friedman, W., McTiernan, R., Yachnis, A., Hayes, D.N., Perou, C.M., Zheng, S., Vegesna, R., Mao, Y., Akbani, R., Aldape, K., Bogler, O., Fuller, G.N., Liu, W., Liu, Y., Lu, Y., Mills, G., Protopopov, A., Ren, X., Sun, Y., Wu, C.J., Yung, W.A., Zhang, W., Zhang, J., Chen, K., Weinstein, J.N., Chin, L., Verhaak, R.G., Noushmehr, H., Weisenberger, D.J., Bootwalla, M.S., Lai, P.H., Triche, T.J., Berg, D.J.V.D., Laird, P.W., Gutmann, D.H., Lehman, N.L., VanMeir, E.G., Brat, D., Olson, J.J., Mastrogianakis, G.M., Devi, N.S., Zhang, Z., Bigner, D., Lipp, E., McLendon, R.: The somatic genomic landscape of glioblastoma. Cell 155(2), 462 – 477 (2013). DOI https://doi.org/10.1016/j.cell.2013.09.034. URL http://www.sciencedirect.com/science/article/pii/S0092867413012087

[5] Corwin, D., Holdsworth, C., Rockne, R.C., Trister, A.D., Mrugala, M.M., Rockhill, J.K., Stewart, R.D., Phillips, M., Swanson, K.R.: Toward patient-specific, biologically optimized radiation therapy plans for the treatment of glioblastoma. PloS one 8(11), e79,115 (2013)

[6] Hamer, P.C.: Small molecule kinase inhibitors in glioblastoma: a systematic review of clinical studies. Neuro-oncology 12(3), 304–316 (2010). DOI 10.1093/neuonc/nop068

[7] Ene, C.I., Holland, E.C.: Personalized medicine for gliomas. Surgical neurology international 6(Suppl 1), S89 (2015)

[8] Eskilsson, E., Røsland, G.V., Solecki, G., Wang, Q., Harter, P.N., Graziani, G., Verhaak, R.G.W., Winkler, F., Bjerkvig, R., Miletic, H.: Egfr heterogeneity and implications for therapeutic intervention in glioblastoma. Neuro-Oncology 20(6), 743–752 (2017). DOI 10.1093/neuonc/nox191. URL http://dx.doi.org/10.1093/neuonc/nox191

[9] Giannini, C., Sarkaria, J.N., Saito, A., Uhm, J.H., Galanis, E., Carlson, B.L., Schroeder, M.A., James, C.D.: Patient tumor egfr and pdgfra gene amplifications retained in an invasive intracranial xenograft model of glioblastoma multiforme. Neuro-Oncology 7(2), 164–176 (2005). DOI 10.1215/s1152851704000821

[10] Giese, A., Bjerkvig, R., Berens, M., Westphal, M.: Cost of migration: invasion of malignant gliomas and implications for treatment. Journal of clinical oncology 21(8), 1624–1636 (2003)

[11] de Groot, J.F., Gilbert, M., Aldape, K., Hess, K.R., Hanna, T., Ictech, S., Groves, M., Conrad, C., Colman, H., Puduvalli, V., et al.: Phase ii study of carboplatin and erlotinib (tarceva, osi-774) in patients with recurrent glioblastoma. Journal of neuro-oncology 90(1), 89–97 (2008). DOI 10.1007/s11060-008-9637-y

[12] Hartung, N., Mollard, S., Barbolosi, D., Benabdallah, A., Chapuisat, G., Henry, G., Giacometti, S., Iliadis, A., Ciccolini, J., Faivre, C., et al.: Mathematical modeling of tumor growth and metastatic spreading: Validation in tumor-bearing mice. Cancer Research 74(22), 6397–6407 (2014). DOI 10.1158/0008-5472.can-14-0721. URL http://dx.doi.org/10.1158/0008-5472.CAN-14-0721

[13] Joo, K.M., Kim, J., Jin, J., Kim, M., Seol, H.J., Muradov, J., Yang, H., Choi, Y.L., Park, W.Y., Kong, D.S., et al.: Patient-specific orthotopic glioblastoma xenograft models recapitulate the histopathology and biology of human glioblastomas in situ. Cell Reports 3(1), 260–273 (2013). DOI 10.1016/j.celrep.2012.12.013. URL http://dx.doi.org/10.1016/j.celrep.2012.12.013

[14] Marin, B., Mladek, A., Burgenske, D., He, L., Hu, Z., Bakken, K., Carlson, B., Schroeder, M., Sarkaria, J.: Ddis-01. the antibody-drug conjugate abt-414 demonstrates single-agent anti-cancer activity across a panel of gbm patient-derived xenografts. Neuro-Oncology 20(suppl-6), vi69–vi69 (2018). DOI 10.1093/neuonc/noy148.280. URL https://doi.org/10.1093/neuonc/noy148.280

[15] Nagane, M., Coufal, F., Lin, H., Böogler, O., Cavenee, W.K., Huang, H.J.: A common mutant epidermal growth factor receptor confers enhanced tumorigenicity on human glioblastoma cells by increasing proliferation and reducing apoptosis. Cancer Res; Cancer Research 56(21), 5079–5086 (1996)

[16] Ningaraj, N.S.: Drug delivery to brain tumours: challenges and progress. Expert opinion on drug delivery 3(4), 499–509 (2006)

[17] Reardon, D.A., Desjardins, A., Vredenburgh, J.J., Gururangan, S., Friedman, A.H., Herndon, J.E., Marcello, J., Norfleet, J.A., McLendon, R.E., Sampson, J.H., et al.: Phase 2 trial of erlotinib plus sirolimus in adults with recurrent glioblastoma. Journal of neuro-oncology 96(2), 219–230 (2010). DOI 10.1007/s11060-009-9950-0

[18] Reardon, D.A., Lassman, A.B., van den Bent, M., Kumthekar, P., Merrell, R., Scott, A.M., Fichtel, L., Sulman, E.P., Gomez, E., Fischer, J., et al.: Efficacy and safety results of abt-414 in combination with radiation and temozolomide in newly diagnosed glioblastoma. Neuro-Oncology 19(7), 965–975 (2017). DOI 10.1093/neuonc/now257. URL http://dx.doi.org/10.1093/neuonc/now257

[19] Van Tellingen, O., Yetkin-Arik, B., De Gooijer, M., Wesseling, P., Wurdinger, T., De Vries, H.: Overcoming the blood–brain tumor barrier for effective glioblastoma treatment. Drug Resistance Updates 19, 1–12 (2015)

[20] Wen, P.Y., Kesari, S.: Malignant gliomas in adults. New England Journal of Medicine 359(5), 492–507 (2008)

